# Loss of mitochondrial co-chaperone GRPEL2 protects mice from age- and diet-induced obesity

**DOI:** 10.64898/2026.05.07.723644

**Authors:** Yang Yang, Nirajan Neupane, Jouni Kvist, Jonna Saarimäki-Vire, Matthias Schewe, Kalle Luopajärvi, Pooja Manjunath, Svetlana Konovalova, Rubén Torregrosa-Muñumer, Veijo Kinnunen, Pekka Katajisto, Timo Otonkoski, Eija Pirinen, Jayasimman Rajendran, Henna Tyynismaa

## Abstract

Mitochondrial protein homeostasis intersects with metabolic control, but the *in vivo* roles of specific mitochondrial co-chaperones remain unclear. The chaperone mtHSP70 plays a key role in import and folding of nuclear-encoded proteins targeted to mitochondrial matrix. Its protein folding cycle is regulated by the GrpE-like nucleotide exchange factor GRPEL1. Vertebrates also have a GRPEL2 paralog, postulated as the stress-sensitive counterpart, but its physiological relevance is not known. We show here that GRPEL2 is not essential for viability in mice, and its absence does not induce proteotoxic stress responses in stark contrast to GRPEL1. However, we find that GRPEL2 has a role in regulating body weight homeostasis. GRPEL2 knockout mice are protected from age- and diet-induced weight gain and maintain a better metabolic health and insulin sensitivity. Transcriptional profiling revealed minimal changes in liver and skeletal muscle, whereas white adipose tissue from Grpel2-deficient mice lacked the obesity-associated remodeling seen in controls. We propose that GRPEL2 fine-tunes metabolic setpoints without broadly perturbing mitochondrial protein import, thereby maintaining adipose tissue health during nutritional excess. These findings show that subtle alterations in mitochondrial chaperone systems reshape systemic metabolism and could suggest strategies to mitigate obesity and insulin resistance through targeted modulation of mitochondrial proteostasis.

## Introduction

Mitochondria contribute to whole-body energy homeostasis through multiple different mechanisms. Mitochondrial respiratory chain (RC) function is vital for meeting energy demands, but mitochondria also host central biosynthetic processes such as those contributing to folate, nucleotide and iron-sulfur cluster (ISC) synthesis (Friedman & Nunnari, 2014; Lill & Freibert, 2020; Suomalainen & Nunnari, 2024). The tissue-specific manifestation of primary mitochondrial diseases suggests that a range of factors are involved in modifying mitochondrial functions to meet the specific requirements of each tissue (Durieux *et al*, 2011; Ott *et al*, 2016; Pagliarini *et al*, 2008; Suomalainen & Battersby, 2018).Many of the adaptative regulators of mitochondrial metabolism are not yet well understood.

Critical for mitochondrial functions is its proteome, which is largely encoded by nuclear genes, synthesized by cytosolic ribosomes, and transported into mitochondria through specialized import pathways. Mitochondrial matrix proteins are transferred through the double membrane by outer and inner membrane translocase complexes (TOM and TIM) (Chacinska *et al*, 2009; Chacinska *et al*, 2005; Milenkovic *et al*, 2009; Model *et al*, 2001; Neupert & Herrmann, 2007; Saitoh *et al*, 2007; Wiedemann & Pfanner, 2017). On the matrix side, the presequence translocase-associated import motor (PAM) complex assists in directing the imported polypeptides into the matrix. The mitochondrial chaperone mtHSP70 forms the core of the import motor and initiates the folding of the entering polypeptide (Horst *et al*, 1997; Kampinga & Craig, 2010; Kang *et al*, 1990; Krayl *et al*, 2007; Liu *et al*, 2003; van der Laan *et al*, 2007). The client protein binding and release cycle of mtHSP70 is controlled by J-proteins, which regulate client capture by accelerating ATP hydrolysis, and by GrpE-like nucleotide exchange factors (NEF), which stimulate the release of ADP to prepare the mtHSP70 for another cycle of protein folding (Chacinska *et al*., 2009; Kampinga & Craig, 2010; Liu *et al*., 2003; Slutsky-Leiderman *et al*, 2007).

Two mitochondrial GrpE-like NEFs, GRPEL1 and GRPEL2, are known to exist in vertebrates (Naylor *et al*, 1998) and both are ubiquitously expressed. Recent studies have shown that GRPEL1 is the canonical NEF in mammalian mitochondria (Konovalova *et al*, 2018; Manjunath *et al*, 2024; Morizono *et al*, 2024; Morizono *et al*, 2026; Srivastava *et al*, 2017). In support of its essential role in mammals, GRPEL1 knockout mice are embryonal lethal (Neupane *et al*, 2022). Furthermore, inducible depletion of GRPEL1 in mouse skeletal muscle resulted in rapid muscle atrophy with shutdown of mitochondrial metabolism and pronounced induction of proteotoxic stress responses as a result of protein mistargeting and misfolding (Neupane *et al*., 2022). On the contrary, cultured human cells lacking GRPEL2 are viable and able to import proteins into mitochondria (Konovalova *et al*., 2018). GRPEL2 dimerization into its presumed active form is redox-regulated, and it may thus act as a stress-adaptive fine-tuner of mitochondrial proteostasis (Konovalova *et al*., 2018; Morizono *et al*., 2026). To investigate the physiological relevance of GRPEL2, we generated whole-body knockout mice. Our results show that GRPEL2 is not essential for viability but it modifies the body weight homeostasis of adult mice with beneficial effects on metabolic health. Our findings suggest that GRPEL2 has gained a specialized role as a metabolic regulator in mammals.

## Results

### Mice lacking GRPEL2 are viable and resistant to age-related weight gain

*Grpel2* knockout (*Grpel2* KO) mice were born at Mendelian ratios and appeared healthy (Fig 1A, B). We confirmed the absence of *Grpel2* on mRNA and protein levels (Fig S1A, B). We monitored the weight of *Grpel2* KO mice until one year of age and observed that although their food intake was comparable to WT C57Bl/6 littermates (Fig 1C), *Grpel2* KO mice of both genders had a progressively lower body weight gain (Fig 1D). The body weight difference in comparison to WT littermates was 19% for males and 23% for females at 47 weeks of age. Follow-up in metabolic cages showed that *Grpel2* KO mice and WT littermates were equally active (Fig 1E). Body composition analysis at one year of age indicated overall lower fat and lean mass in *Grpel2* KO mice but no differences when calculated in proportion to total body weight (Fig 1F, Fig S1C). Similarly, organ weights were lower in *Grpel2* KO but organ to body weight ratios were unchanged (Fig 1G, Fig S1D-E). These results indicated that *Grpel2* KO mice were proportionally slightly smaller than their WT littermates. By indirect calorimetry, we observed that *Grpel2* KO mice consumed more oxygen than their WT littermates (Fig 1H). Energy expenditure was not different between WT and KO mice after adjusting by ANCOVA as recommended by (Tschop *et al*, 2011) (Fig 1I). Respiratory exchange ratio (RER) indicating substrate utilization was not altered (Fig 1J).

**Figure 1.**
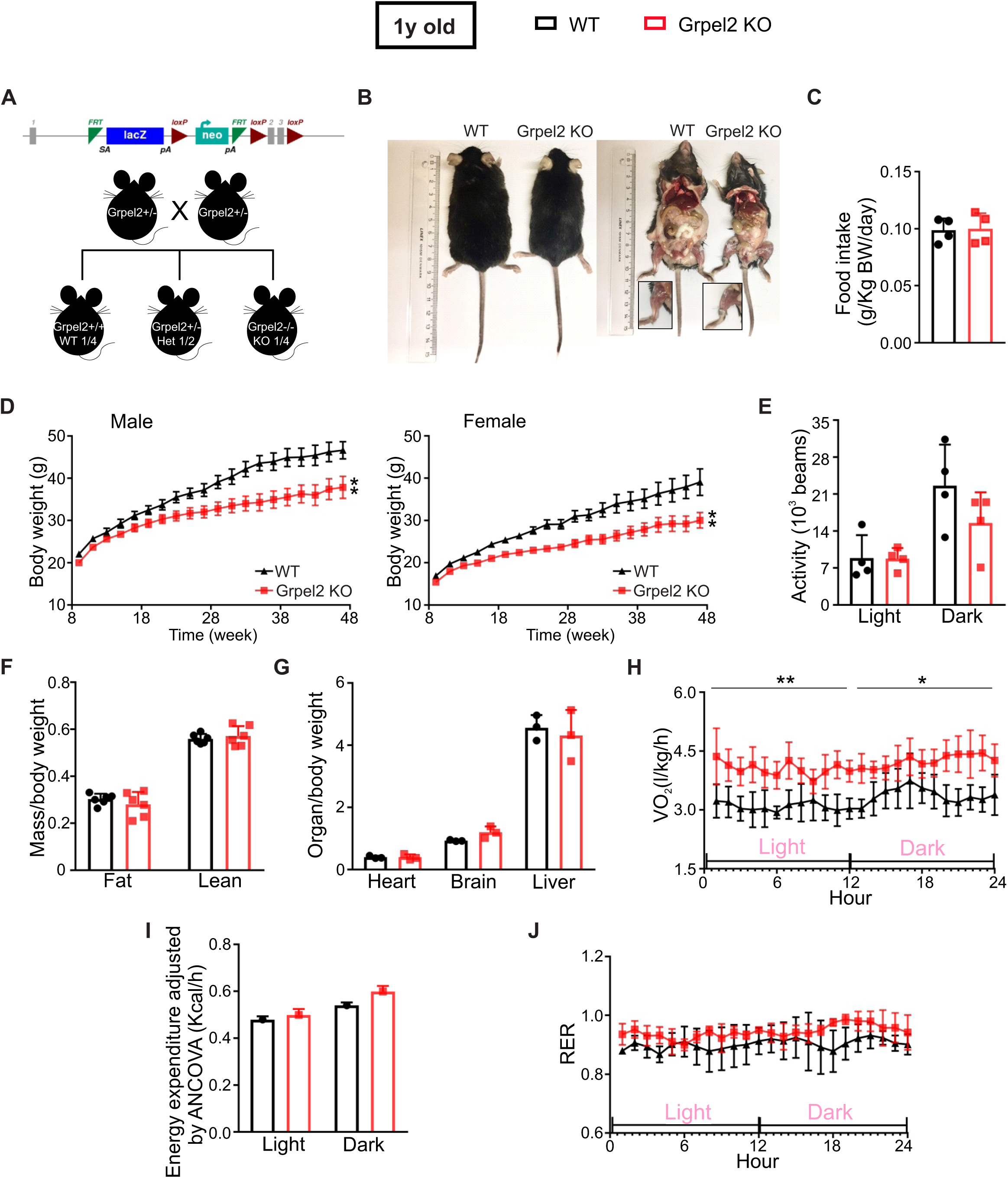
Grpel2 KO mice are viable but gain progressively less weight. (A) Breeding scheme for generating Grpel2 KO mice. (B) Appearance of one-year-old WT and Grpel2 KO mice. (C) Daily food intake (n = 4/genotype). (D) Body weight gain of male and female mice (n = 8-10/genotype) (E) Spontaneous activity of mice (n = 4/group). (F) Fat and lean mass normalized to body weight (n = 6/group). (G) Tissue weight of heart, brain and liver normalized to body weight (n = 3/group). (H) Oxygen consumption normalized to body weight as measured by indirect calorimetry (n = 4-6/genotype). (I) Energy expenditure adjusted by ANCOVA. (J) Respiratory exchange ratio normalized to body weight (n = 4-6/genotype). Data in B-K were collected from one-year-old mice. All data are shown as mean, and error bars indicate SD. ∗p ≤ 0.05, ∗∗ p ≤ 0.01.

The C57BL/6 mouse strain is widely used in metabolic and obesity research because of its susceptibility to age-related obesity and insulin resistance in *ad libitum* fed conditions. We performed glucose (GTT) and insulin tolerance (ITT) tests on the one-year-old mice and found that *Grpel2* KO mice exhibited better glucose clearance capability (Fig 2A) and insulin sensitivity (Fig 2B) than WT littermates. Insulin levels and pancreatic beta cell mass were not different between WT and *Grpel2* KO mice (Fig S2), suggesting that increased insulin secretion was not the primary mechanism behind the superior glucose tolerance of the KO mice. Hematoxylin and eosin (HE) staining showed reduced hepatocellular vacuolation in the *Grpel2* KO liver (Fig 2C) and Oil Red O staining indicated less fat accumulation (Fig 2D). HE staining on epididymal white adipose tissue (eWAT) displayed smaller and more uniformly organized adipocytes in *Grpel2* KO mice than in WT littermates (Fig 2E, F). Intestinal crypts had normal appearance supporting that the smaller size and poorer weight gain of the *Grpel2* KO mice were not caused by defects in food absorption (Fig S3A). Total cholesterol and LDL were significantly lower in the blood serum of one year old *Grpel2* KO mice whereas triglycerides, total NEFA (Non-Esterified Fatty Acids), and HDL were unchanged (Fig S3B-F). Thus, *Grpel2* KO mice were resistant to age-associated weight gain and maintained a widely better metabolic health than their littermates.

**Figure 2.**
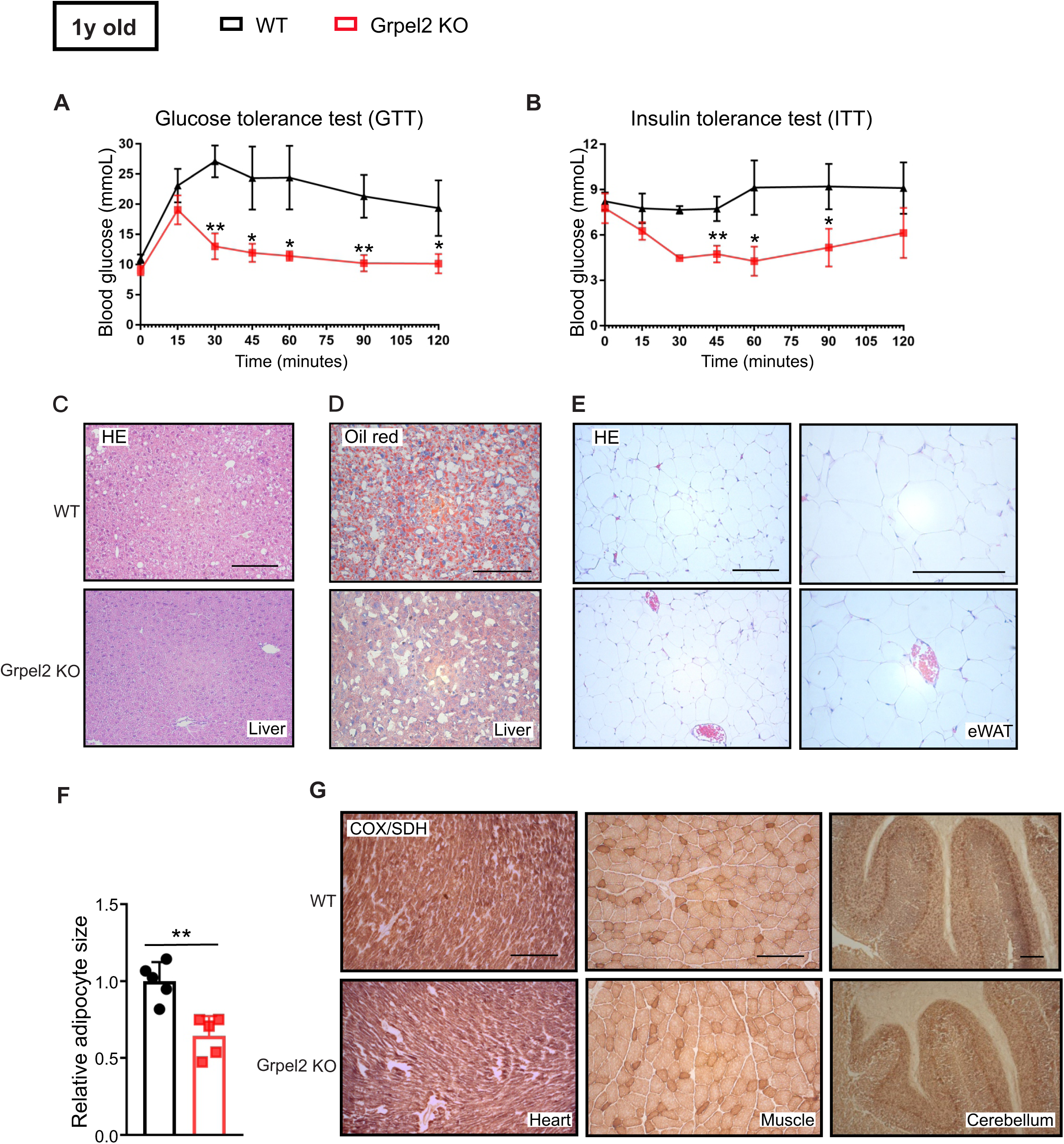
One-year-old Grpel2 KO mice are resistant to age-related metabolic alterations. (A) Glucose tolerance test (n = 5-6/genotype). (B) Insulin tolerance test (n = 5-6/genotype). (C) HE staining of liver. Scale bars, 100 μm. (D) Oil Red staining of liver showing hepatic lipid content. Scale bars, 100 μm. (E) HE staining of eWAT. Scale bars, 50 μm, 100 μm. (F) Quantification of adipocyte size (n = 5 mice/genotype), 5 slides from each mouse were analyzed with 8-10 images from each slide randomly chosen for quantification. (G) Histochemical staining of cytochrome c oxidase (COX; brown) and succinate dehydrogenase (SDH; blue) activities of heart, muscle and cerebellum tissues. Scale bars, 100 μm. All data were collected from one-year-old mice. Graphs show the data as mean, and error bars indicate SD. ∗p ≤ 0.05, ∗∗ p ≤ 0.01.

Loss of a critical mitochondrial protein may disrupt RC function, which becomes evident in mitochondria-rich tissues such as the heart, skeletal muscle, and brain. However, we did not detect clear signs of RC deficiency in these tissues in *Grpel2* KO mice. Even at the age of one year, stainings for RC complex II (SDH) and IV (COX) activities were normal (Fig 2G).

### Transcriptome analysis indicates lack of obesity-induced eWAT remodeling in *Grpel2* knockout mice

To gain understanding of how the absence of GRPEL2 affected the metabolism of the KO mice, we performed RNA sequencing on the skeletal muscle, liver, and eWAT of one-year-old *Grpel2* KO mice and their WT littermates. Principal component analysis revealed that the transcriptomes of liver and muscle in *Grpel2* KO mice were largely indistinguishable from those of WT littermates, whereas eWAT transcriptome differed widely between the genotypes (Fig 3A-C). The number of differentially expressed genes (FDR≤0.01) was 5 for skeletal muscle, 43 for liver and 2903 for eWAT. Downregulation of *Grpel2* was the only alteration common to all three tissues (Fig 3D, Table S1). Expression levels of *Hspa9* (mtHSP70) or *Grpel1* were not changed (Fig 3E).

**Figure 3.**
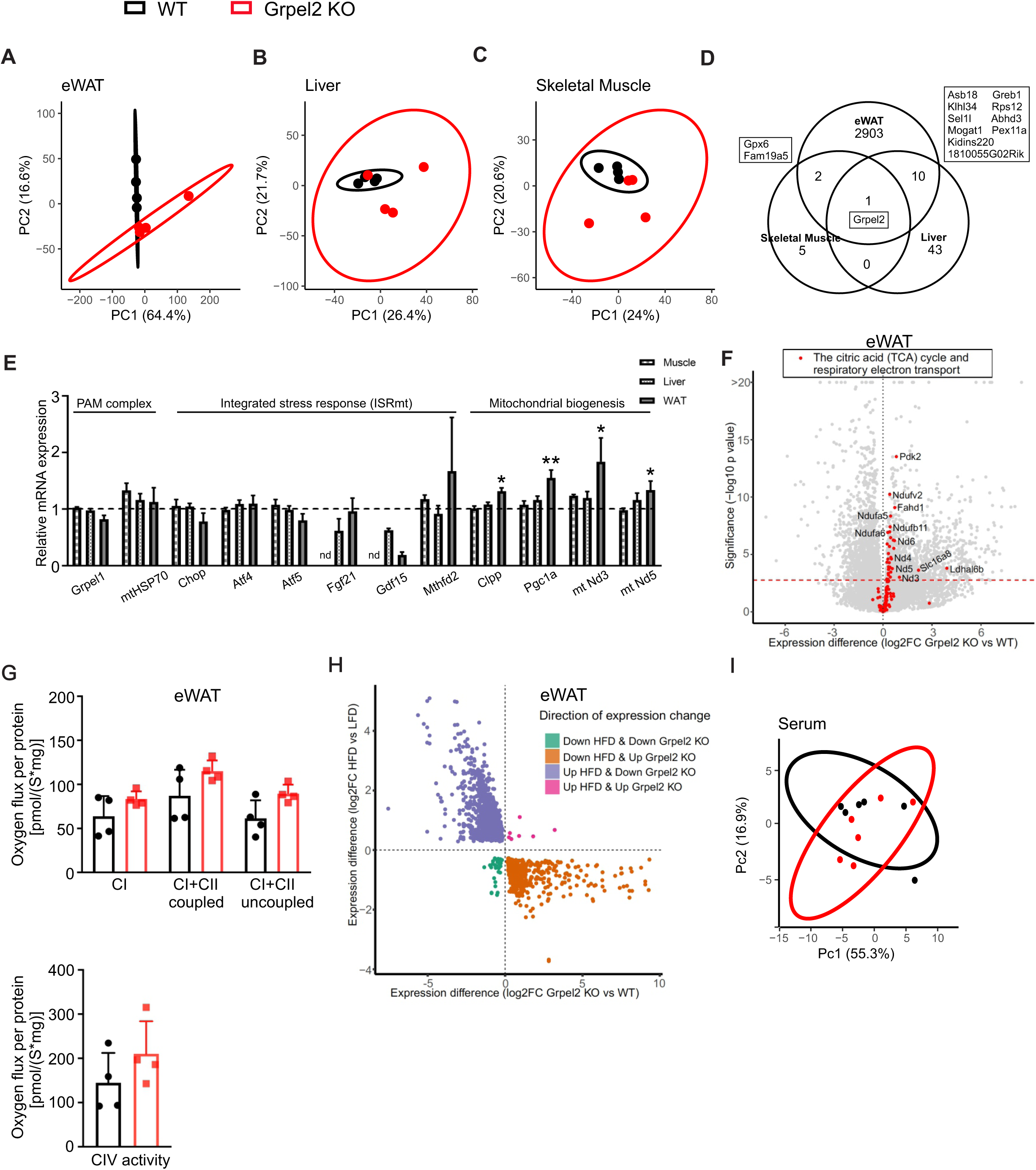
Lack of obesity-related remodeling in Grpel2 KO eWAT. (A-C) Principal component analysis for RNA sequencing data from eWAT, liver and skeletal muscles of one-year-old mice. The ellipses around the samples indicate the 95% confidence intervals (n=4/genotype). (D) Principal component analysis for targeted metabolomics data from serum of one-year-old mice (n=6/genotype). (E) Venn Diagram of differentially expressed genes (FDR<0.01) in adipose tissue, liver and skeletal muscle. (F) Gene expression of selected genes in PAM complex, integrated stress response and mitochondrial biogenesis pathways. (G) Respirometry analyses of coupled respiration through either Complex I or Complex I+II in isolated mitochondria from eWAT (n = 4/genotype). Respirometry analyses of Complex IV activity in isolated mitochondria from eWAT (n = 4/genotype). (H) Volcano plot of eWAT genes, highlighting altered genes in the TCA cycle and electron transport pathway in red. The statistically significant expression changes (FDR<0.01) are marked by a dashed red line. (I) Scatterplot of expression differences in white adipose tissue resulting from alteration in diet (high fat diet (HFD) vs low fat diet (LFD); dataset GSE63198) and Grpel2 KO vs WT (this study). Results indicate opposite changes in Grpel2 KO mice as compared to the effects of HFD.

Dysregulated mitochondrial proteostasis induces transcriptional stress responses such as mitochondrial unfolded protein response (UPR) or integrated stress response (Anderson & Haynes, 2020; Forsstrom *et al*, 2019; Quiros *et al*, 2017), which were pronounced in the skeletal muscle specific *Grpel1* KO mice (Neupane *et al*., 2022). However, in the studied *Grpel2* KO tissues we did not detect an induction of stress markers *Chop*, *Atf4*, *Atf5*, *Fgf21*, *Gdf15* or *Mthfd2* (Fig 3E). Mitochondrial biogenesis markers such as *Clpp*, *Ppargc1a* (Pgc1a), and nuclear- and mtDNA-encoded genes for respiratory chain subunits were somewhat increased in eWAT, but not in muscle or liver (Fig 3E, F). The higher level of mitochondrial biogenesis transcripts in *Grpel2* KO eWAT is not likely to indicate a compensatory response to dysfunctional mitochondria, but represents the higher mitochondrial content in the leaner knockout eWAT in comparison to their obese WT littermates (Heinonen *et al*, 2015; Heinonen *et al*, 2017; Lin *et al*, 2016). Indeed, when we measured the oxygen flux in isolated eWAT mitochondria using a high-resolution respirometry we did not observe significant differences between *Grpel2* KO and WT mice (Fig 3G).

We noticed that the majority of the differentially expressed genes and pathways that were altered between *Grpel2* KO and WT littermate eWAT were the same as those differing between C57BL/6 mouse eWAT on low and high fat diets (GSE63198, (Choi *et al*, 2015)), but to the opposite direction (Fig 3H, S4B). Besides genes on mitochondrial pathways, these included genes related to inflammatory responses (chemokines *Ccl2*, *Ccl3*), extracellular matrix components such as collagens, cathepsins, matrix metalloproteinases, and genes related to antioxidant defense (*Gpx3*), fatty acid uptake (*Fabp4*, *Cd36*, *Slc27a2*), elongation (*Elovl6*), or activation (*Acsm3*, *Acacb*, *Acot4*, *Acadsb*, *Hadh*, *Faah*). No markers for eWAT browning were detected as elevated in *Grpel2* KO mice. Overall, the transcriptome findings suggested that the absence of GRPEL2 had little effect in skeletal muscle and liver, even at one year of age. The major alterations observed in eWAT were likely to be secondary to the ability of *Grpel2* KO mice to resist excessive weight gain.

We also performed a targeted metabolite profiling in serum samples of one-year-old mice, which showed comparable profiles in *Grpel2* KO and WT littermates (Fig 3I). Serum ornithine, asparagine, choline, acetoacetic acid and adenine levels were altered in *Grpel2* KO mice (Fig S4A), but not significantly after correcting for multiple testing.

### *Grpel2* knockout mice are resistant to high fat diet induced weight gain

As *Grpel2* KO mice were resistant to the age-related weight gain, which is a characteristic feature of C57/Bl6 strain, we next investigated if they could also resist diet-induced obesity at young age. WT and *Grpel2* KO mice were fed either with a high-fat diet (HFD) containing 60% calories from fat or a control diet (CD) containing 6% calories from fat, for ten weeks starting at the age of 8 weeks (Fig 4A). The HFD induced considerable weight gain in WT mice but not in *Grpel2* KO mice (Fig 4B-D). Both WT and *Grpel2* KO mice on CD gained approximately 30% of body weight as they grew in size from the age of 8 to 18 weeks. However, on HFD, WT mice gained approximately 60% of body weight whereas *Grpel2* KO mice gained no more than the 30% corresponding to the growth on CD (Fig 4E). WT and *Grpel2* KO mice consumed comparable amounts of HFD food per body weight (Fig 4F). Ratios of fat or lean mass to body weight were not different between genotypes after CD (Fig 4G), but following HFD, *Grpel2* KO mice had less fat mass than WT mice (Fig 4H). We measured oxygen consumption and found that the young *Grpel2* KO mice on CD consumed the same amount of oxygen as WT mice, whereas on HFD *Grpel2* KO mice consumed more oxygen than their WT littermates (Fig 4I). RER curves indicated that HFD altered substrate utilization similarly in both genotypes from carbohydrates to fat (Fig 4J). Activity levels or energy expenditure adjusted by ANCOVA were not altered in *Grpel2* KO mice under either diet (Fig 4K, L).

**Figure 4.**
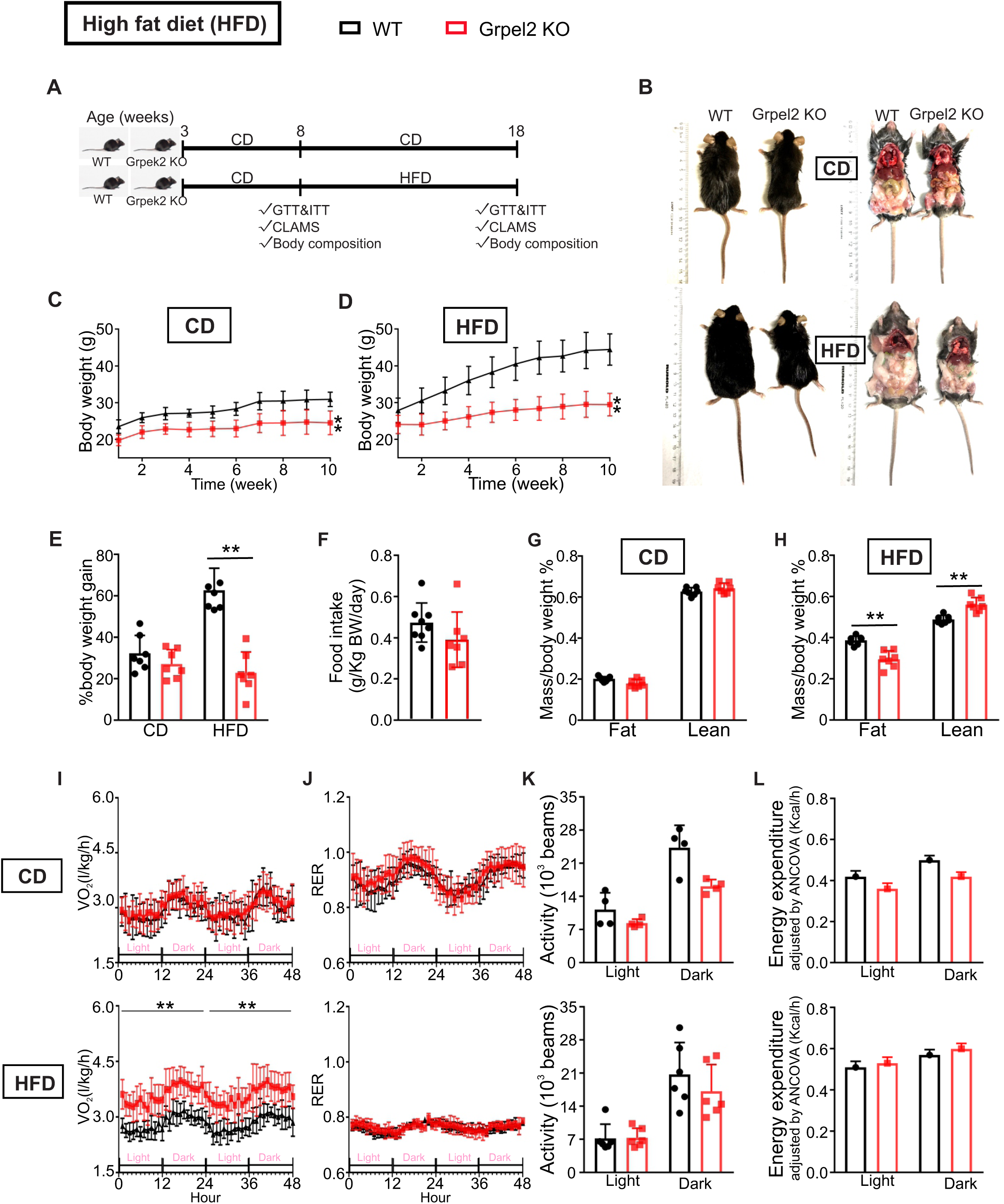
Grpel2 KO mice are resistant to excessive weight gain on high fat diet. (A) Scheme for ten-week high fat diet (HFD) and control diet (CD) experiment. (B) Representative images of WT and Grpel2 KO mice fed with CD or HFD for 10 weeks. (C) Body weight gain on CD (n = 7-8/genotype). (D) Body weight gain on HFD (n = 7-8/genotype). (E) Percentage of body weight gain % at the end of diet (n = 7-8/group). (F) Quantification of daily HFD food intake (n = 7-8/genotype). (G) Fat and lean mass normalized to body weight after CD (n = 6-7/genotype). (H) Fat and lean mass normalized to body weight after HFD (n = 6-7/genotype). (I-L) Oxygen consumption, respiratory exchange ratio (RER), spontaneous activity and energy expenditure in CLAMS after CD and HFD (n = 4-6/group). All data are shown as mean, and error bars indicate SD. ∗p ≤ 0.05, ∗∗ p ≤ 0.01.

Histopathological screening revealed that the absence of GRPEL2 protected the liver (Fig 5A, B) and eWAT from HFD-induced fat accumulation (Fig 5C, D). The eWAT of *Grpel2* KO mice on HFD had small and organized adipocytes in comparison to WT littermates (Fig 5C-E). We performed GTT and ITT on mice before (at 8 weeks of age) and after the 10-week diets. Young *Grpel2* KO mice before the diet and after CD had nearly normal GTT and ITT. However, HFD induced glucose intolerance and insulin resistance in WT mice, which were not observed in *Grpel2* KO mice (Fig 5F, G). Also, serum cholesterol, HDL, and LDL levels were lower in *Grpel2* KO mice than in WT on HFD (Fig S3B-F). These results were consistent with the ability of *Grpel2* KO mice to resist HFD-induced obesity and the consequent metabolic defects.

**Figure 5.**
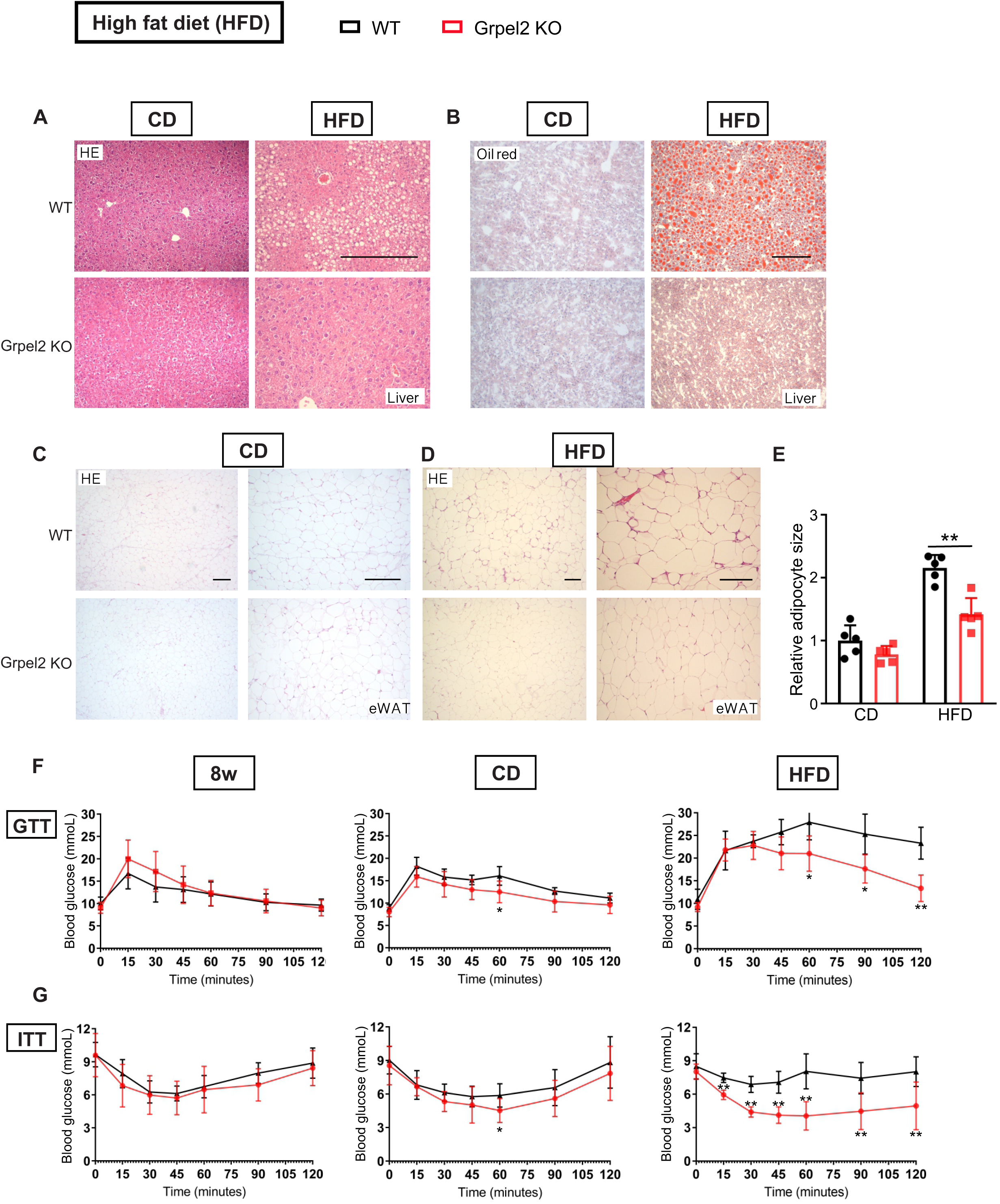
Grpel2 KO mice are resistant to diet-induced metabolic alterations. (A) Representative images of HE stainings of liver after CD or HFD. Scale bars, 100 um. (B) Representative images of Oil red O staining showing fat content in liver after CD or HFD. Scale bars, 100 um. (C-D) Representative images of HE stainings of eWAT after CD or HFD. Scale bars, 100 um. (E) Quantification of adipocyte size (n = 5 mice/genotype), 5 slides from each mouse were analyzed with 8-10 images from each slide randomly chosen for quantification. (F-G) Glucose tolerance test and insulin tolerance test before and after CD or HFD (n = 5-6/group). All data are shown as mean, and error bars indicate SD. ∗p ≤ 0.05, ∗∗ p ≤ 0.01.

## Discussion

Here we identify an *in vivo* role for the mitochondrial co-chaperone GRPEL2 in metabolic regulation. This co-chaperone of mtHSP70 is not essential for life, but has an impact on body weight homeostasis. Absence of GRPEL2 in mice largely prevented age- and diet-induced weight gain and the consequent metabolic defects including insulin resistance. In contrast to GRPEL1, which is the primary NEF regulating the protein folding cycle of mtHSP70 and protein import into mitochondrial matrix, the absence of GRPEL2 in mice did not induce proteotoxic stress responses that typically result from accumulation of mistargeted and misfolded mitochondrial proteins in models of defective mitochondrial protein import (Wrobel *et al*, 2015). These findings indicate that GRPEL2 plays little, if any, role in mitochondrial protein import or proteostasis, at least in the tissues studied here, but it may have instead evolved as a fine-tuning metabolic regulator in mammals.

Few links between GRPEL2 and metabolic regulation have been previously suggested. In a mouse model with acute fatty liver disease, which was induced by blocking long-chain fatty acid beta-oxidation, *Grpel2* expression was reported increased (van der Leij *et al*, 2007). In contrast, another study found that liver and skeletal muscle of mice with reduced insulin gene dosage had *Grpel2* as the only consistent and significantly downregulated gene across diets (Templeman *et al*, 2017). In addition, in different transcriptome datasets, liver *Grpel2* was consistently decreased by caloric restriction and increased in a number of HFD studies, as well as in obese ob/ob mice (Templeman *et al*., 2017). These patterns raise the possibility that GRPEL2 contributes to metabolic adaptation to fuel availability. Speculatively, the evolutionary acquisition of GRPEL2 may have supported energy conservation under limited food intake.

Severe mitochondrial respiratory dysfunction results in many disease conditions while mild perturbation can be beneficial. For instance, the widely used anti-diabetic drug metformin is known to induce mild mitochondrial dysfunction and also weight loss (Coll *et al,* 2020). In mice, mild decrease of RC function can be beneficial for metabolic health. For example in the adipose specific knockout of *Tfam*, the mitochondrial transcription factor A, the moderate complex I and IV deficiency resulted in a shift from complex I to complex II dependent respiration and increased beta-oxidation (Vernochet *et al*, 2012). Also the full body knockout of mitochondrial protease *Clpp* protected mice from diet-induced obesity and insulin resistance (Becker *et al*, 2018; Bhaskaran *et al*, 2018). Furthermore, drug-induced inhibition of mitochondrial transcription has been reported to induce metabolic rewiring that reverses diet-induced obesity and hepatosteatosis (Jiang *et al*, 2024). Our findings are consistent with GRPEL2 acting as another fine-tuner of mitochondrial function with impact on body weight maintenance and metabolic health, through currently unknown mechanism.

What then could be the molecular function of GRPEL2 by which it contributes to metabolic regulation? *In vitro*, GRPEL2 is capable of functioning as NEF for mtHSP70 (Srivastava *et al*., 2017), which supports its primary role in protein folding. Proximity labeling (BioID) identified mtHSP70, GRPEL1, and HSPE1 as proximal partners, situating GRPEL2 near the mitochondrial import and protein folding machinery (Konovalova *et al*., 2018). GRPEL2 was also proximal to enzymes in the 2-oxoglutarate dehydrogenase and branched-chain α-keto acid dehydrogenase complexes, pointing to potential roles at the interface of protein quality control and central carbon/amino acid metabolism. GRPEL2 has a redox-sensitive cysteine capable of modulating its binding to mtHsp70, which supports the possibility that GRPEL2 has a specific role in oxidative stress (Konovalova *et al*., 2018; Manjunath *et al*., 2024). Under oxidizing conditions, GRPEL2 could modulate selective import or folding of specific substrates, possibly stabilizing mtHSP70-client interactions, reducing premature release and reducing aggregation (Morizono *et al*., 2026). Alternatively, GRPEL2 could support folding or maintenance of a subset of metabolic enzymes, leading to subtle shifts in respiratory chain function or substrate utilization. We note that while *Grpel1* was not upregulated in Grpel2 KO tissues, conversely, *Grpel2* was induced in the skeletal muscle of Grpel1 KO mice (Neupane et al., 2022), suggesting that GRPEL2 may participate in proteotoxic stress responses but cannot compensate for the essential import role of GRPEL1. Together, the data argue for a contextual, possibly tissue-dependent role of GRPEL2, subordinate to GRPEL1 for import but influential for metabolic tuning.

In conclusion, we have shown that mitochondrial co-chaperone GRPEL2 is dispensable for life but contributes to metabolic regulation and body weight homeostasis in mice. Our study in mice did not pinpoint the exact mechanistic role for GRPEL2 in mitochondria, which requires further detailed investigations. Defining GRPEL2-dependent substrates and redox regulation may clarify how subtle alterations in mitochondrial chaperone systems reshape systemic metabolism and could suggest strategies to mitigate obesity and insulin resistance through targeted modulation of mitochondrial proteostasis.

## Methods and materials

### Knockout mice

Embryonic stem (ES) cell line with KO first alleles targeted for *Grpel2* (Grpel2^tm1a(EUCOMM)Hmgu^) were obtained from European Mouse Mutant Cell Repository (EuMMCR). ES cells were expanded and then injected into the blastocyst of C57BL/6J mice in Biocenter Oulu Transgenic Core Facility. The chimeras were genotyped and bred to generate the respective colonies. *Grpel2* KO mice were then crossed with PGK-Cre (Lallemand *et al*, 1998) to remove exons 2 and 3. Animal experiments were performed in compliance with the national ethical guidelines set by the European Union and were approved by the National Animal Experiment Board. The ethical practice of handling laboratory animals was strictly followed throughout the procedures. All experiments were done with male mice except the weight monitoring for one year, which was done with females and males. The mice were housed at 22°C with 12 hour light/dark cycles with *ad libitum* access to either standard rodent diet (CD; 6.2% fat, 44.2% carbohydrate, 18.6% protein; Teklad Global 18% Protein Rodent Diet) or high fat diet (HFD; 60% fat, 20% carbohydrate, 20% protein; Research Diets, Cat#D12492) and water. Body weights of WT and Grpel2 KO mice (both males and females) were recorded weekly. Food intake was measured daily for two weeks. Tissues were collected either by snap-frozen in liquid nitrogen and stored at −80°C until used or fixed with 10% formalin for histology purposes.

### Whole-Body Metabolism and body composition

We used Oxymax Comprehensive Lab Animal Monitoring System (CLAMS; Columbus Instruments) to measure oxygen consumption, carbon dioxide production and spontaneous activity of the mice. Mice were in CLAMS for 24-48 hours at room temperature (+22°C). Food and water were available *ad libitum*. Analysis of covariance (ANCOVA) was used for normalization using the NIDDK Mouse Metabolic Phenotyping Center’s (MMPC, www.mmpc.org) Energy Expenditure Analysis (www.mmpc.org/shared/regression.aspx). Bruker minispec LF50 Body Composition Analyser was used to measure the body composition including fat and lean mass (Muller *et al*, 2021).

### Glucose and insulin tolerance tests

For the oral glucose tolerance test, mice were fasted for 6 hours prior to test, and then challenged by oral glucose (1g/kg). Blood glucose levels were measured by tail vein puncture prior to glucose administration, and at 15, 30, 60, 90, and 120 minutes after glucose administration with a portable glucometer (Accu-Chek Aviva Glucose Meter, Roche). For the insulin tolerance test, mice were challenged by i.p. insulin injection of 0.8 U/kg (Novolin R, Novo Nordisk). Blood glucose levels were measured immediately prior to insulin administration, and at 15, 30, 60, 90, and 120 minutes after insulin administration.

### Biochemical analysis of blood serum

Blood samples were collected by cardiac puncture. Total serum cholesterol, HDL, LDL, triglycerides, non-esterified fatty acids and iron were analyzed with a Siemens ADVIA 1650 analyzer (Diamond Diagnostics) at the Biochemical Analysis Core for Experimental Research, University of Helsinki. Plasma insulin levels were determined from tail vein by Ultra Sensitive Mouse Insulin ELISA kit (Crystal Chem) according to the manufacturer’s instructions.

### Histology

Tissue samples were fixed with 10% buffered formalin for overnight at 4°C. After washing with 70% ethanol and preparation procedures, tissues were mounted in paraffin blocks. Paraffin-embedded tissues were sectioned (5um thickness) and stained with HE (Hematoxylin and Eosin stain) followed the standard protocol (Khan *et al*, 2017; Tyynismaa *et al*, 2005). For COX/SDH double staining, tissues were collected in OCT Compound Embedding Medium (Tissue-Tek) and snap-frozen in 2-methylbutane bath in liquid nitrogen (Khan *et al*., 2017; Tyynismaa *et al*., 2005).

Hepatic lipid accumulation was analyzed by Oil Red O staining of frozen liver sections (Ahola-Erkkila *et al*, 2010). Stained sections were analyzed by Axioplan 2 Universal Microscope (Zeiss). The adipocyte sizes were quantified double blindly using Image-J, with 5-10 images from each slide randomly selected throughout the area.

### Immunofluorescence

Tissues were fixed in 4 % PFA and embedded in paraffin. Paraffin blocks were cut into 5 µm sections with a microtome. After deparaffinization the antigen retrieval was done by boiling slides either in 1 mM EDTA or 0.1 M citrate buffer, blocking was performed for 30 min at RT in blocking buffer (2 % BSA, 5 % Normal goat serum, 0.375 % Triton X-100, in PBS). For Olfm4 and Lysozyme, Dako Target Retrieval solution S169984-2 was used. Primary antibodies were diluted in blocking buffer and incubated overnight at +4°C. Secondary antibodies and DAPI were diluted in blocking buffer and incubated 45 min at RT. The samples were mounted with Shandon Immu-Mount mounting media. Images were taken with Leica DM5000B widefield microscope.

### Immunoblots

We prepared protein lysates from 20mg of snap frozen tissues in RIPA buffer (Cell signaling technology) containing Halt™ Protease Inhibitor Cocktail (ThermoFisher). We homogenized the samples using Fast Prep w-24 Lysing Matrix D (MP Biomedical) and Precellys w-24 (Bertin Technologies), as described in (Forsstrom *et al*., 2019). Protein concentrations were quantified with Bradford method (Bio-Rad). We loaded 20 μg of protein per sample in 10% stain free polyacrylamide gels (Bio-Rad) and transferred to PVDF membranes using transfer-blot turbo transfer system (Bio Rad). After blocking with 5% milk in 1X TBS-T and sequent washing, membranes were incubated with primary antibodies overnight at 4°C (Star Method, antibody table). Following secondary antibody incubation and Clarity Western ECL Substrate (BioRad), protein bands were detected by Chemidoc XRS+ Molecular Imager (Bio-Rad). Quantification was done with the ImageLab software (Bio Rad).

### RNA sequencing

Tissue samples were homogenized using Fast Prep w-24 Lysing Matrix D (MP Biomedical) and Precellys w-24 (Bertin Technologies). Total RNA was extracted from snap frozen tissues with miRNAeasy (Qiagen). RNA quality was determined with RNA ScreenTape (Agilent Technologies). All samples had the RNA integrity number (RIN) value greater than 8. NEBNext Ultra II Directional RNA Library Prep Kit for Illumina was used to generate cDNA libraries for next-generation sequencing. 1000 ng of total RNA was used, and poly(A) selection was made with NEBNext Poly(A) mRNA Magnetic Isolation Module. cDNAs were purified with Agencourt AMPure XP beads and were then end-repaired, and the adapter-ligated utilizing dA-tailing. The amplified library was then purified using AMPure XP Beads. Library quality was assessed by TapeStation (Agilent High Sensitivity D5000 assay) and library quantity by Qubit. Prepared libraries were sequenced with Illumina NextSeq500-system (High Output 1 x 75bp paired-end).

Raw data (bcl-files) was demultiplexed, and the adapter sequence (AGATCGGAAGAGCACACGTCTGAACTCCAGTCA) removed with bcl2fastq2 (v2.20.0.422; Illumina), retaining reads with a minimum length of 35bp. The resulting fastq-files were mapped to the mouse reference genome (GRCm38) with STAR (v. 2.5.3; (Dobin *et al*, 2013)). The differential gene expression was analyzed with R (v.3.6.0; (Team, 2019)). The read counts for genes were extracted with Rsubreads (v.1.34.6; (Liao *et al*, 2014)) excluding duplicates, multi-mapping reads, chimeric fragments, and reads with mapping quality below 10. Differential gene expression was analyzed with DESeq2 (v.1.24; (Love *et al*, 2014)), comparing knockout to wild type across all tissues as well as separately for each tissue. Genes with low read depth (<50 reads in total) were excluded. The differentially expressed genes were analyzed for pathway enrichment using the Reactome database (Fabregat *et al*, 2016), with ClusterProfiler (v3.12.0; (Yu *et al*, 2012)) and ReactomePA (v1.28.0; (Yu & He, 2016)).

### Real time quantitative PCR

We used 1ug of total RNA to prepare cDNA using Maxima First Strand cDNA Synthesis Kit (Thermo Fisher). We quantified the gene expression using DyNAmo Flash SYBR Green qPCR Kit (ThermoFisher scientific F-415) according to the manufacturer’s instructions in CFX96^TM^ Real-Time PCR Detection System (Bio-Rad). The gene expression was calculated using ddCT method. The primer sequences are in Star Method, primer table.

### Serum metabolomics

We collected the blood samples from one-year-old mice by cardiac puncture and isolated serum samples by centrifuging at 5000g for 10 min. Targeted metabolomic analysis of 100 metabolites was performed using Waters Acquity ultra performance liquid chromatography (UPLC) and triple-quadrupole mass spectrometry analysis at the Finnish Institute of Molecular Medicine Metabolomics Unit.PCA plots were done the same way for the metabolites and RNA-seq data, both using limma/voom.

### Quantification and statistical analysis

Data are presented as ±SD; P-values less than 0.05 were considered significant. Statistical analyses were performed with GraphPad Prism7.03 (GraphPad Software, Inc.). Student’s t tests (unpaired 2-tailed) or two-way ANOVA were used to compare different groups.

## Data and Code Availability

The accession number for the RNA sequencing data reported in this paper is GEO: GSE 135980. RNASeq and metabolomics data generated from this study are also included in supplementary files (Table S1).

## Acknowledgments

Riitta Lehtinen, Jarkko Ustinov, Laura Mäenpää, Taru Hilander and Mugen Terzioglu are thanked for assistance. Members of Anu Suomalainen-Wartiovaara laboratory are acknowledged for providing protocols and advice. Institute for Molecular Medicine Finland (FIMM) Metabolomics, Electron Microscopy Unit of the Institute of Biotechnology, Biomedicum Functional Genomics Unit, Faculty of Medicine Biochemical analysis core for experimental research and the Histology lab, Finnish Centre for Laboratory Animal Pathology (FCLAP) and Biocenter Oulu Transgenic Core Facility are acknowledged for their services and collaborations. This work was supported by European Research Council (grant number 637458), University of Helsinki, and Sigrid Juselius Foundation to H.T., Research Council of Finland (312438) to P.K., T.O. and H.T., Biomedicum Helsinki Foundation to N.N., Macau University of Science and Technology Faculty Research Grant (FRG-25-059-FMD) and Guangdong Basic and Applied Basic Research Foundation (2021A1515110217) to Y.Y. Part of this work has been supported by the INFRAFRONTIER-I3 project under the EU contract Grant Agreement Number 312325 of the EC FP7 Capacities Specific Programme.

## Supplementary figures and Figure legends

**Figure S1. related to Figure 1. Support data of Grpel2 KO mice characterization**

(A) Immunoblotting of mitochondrial proteins in mouse liver mitochondria extracts. SDHA was used as loading control.

(B) mRNA expression level of *Grpel2* in liver, muscle and eWAT of Grpel2 KO mice (n = 4/genotype).

(C) Fat and lean mass of one-year-old WT and Grpel2 KO mice (n = 6/group).

(D) Tissue weight of heart, brain and liver of one-year-old mice (n = 3/genotype).

(E) Total body weight of the mice in D (n = 3/genotype).

All data are shown as mean, and error bars indicate SD. ∗p ≤ 0.05, ∗∗ p ≤ 0.01.

**Figure S2. related to Figure 2. Basal blood insulin level and pancreatic beta cell characterization**

(A) Basal insulin level in WT and Grpel2 KO mice at one year of age (n = 5-6/genotype).

(B-C) Immunofluorescence staining of insulin (red) and glucagon (green), Hochest stain the nucleus (blue) in WT and Grpel2 KO from CD and HFD groups. Beta cell areas were quantified, and the ratio to the total area are shows in graph. Scale bars, 100 μm.

**Figure S3. related to Figure 2 and Figure 5. Characterization of intestinal crypts and biochemical analysis of serum**

(A) HE staining and immunofluorescence staining of Olfm4 (green) and Lysozyme (red) of ileum samples of WT and Grpel2 KO mice. Scale bars, 100 μm.

(B-F) Graphs showing total cholesterol, triglycerides, total NEFA, HDL and LDL levels in HFD, CD and one-year-old mouse serum (n = 5-6/group).

**Figure S4. related to Figure 3. Targeted serum metabolome analysis and RNAseq pathway analysis of eWAT**

(A) Metabolome analysis of choline, asparagine, ornithine, adenine and acetoacetic acid in one-year-old mice (n= 6/genotype).

Enrichment analysis of differentially expressed genes in adipose tissue, in Grpel2 KO vs WT and HDF vs LDF. Showing the most significantly enriched categories for GO (gene ontology), KEGG (Kyoto Encyclopedia of Genes and Genomes) and Reactome for both dataset. The color indicates the level of significance (red = highly significant, blue =marginally significant) and the size of the circle the levels of enrichment (large circle = high enrichment, small = low enrichment).

**Supplementary data files**

Table S1. (A) Table of differentially expressed genes in Grpel2 KO compared to WT in the eWAT, liver, skeletal muscle of 1y old mice. (B) Table of enrichment results for GO, KEGG and Reactome. The analyses were carried out separately for the up- and down-regulated genes. (C) Metabolomic profiling of 1y old mice analyzed with limma.

## References

Ahola-Erkkila S, Carroll CJ, Peltola-Mjosund K, Tulkki V, Mattila I, Seppanen-Laakso T, Oresic M, Tyynismaa H, Suomalainen A (2010) Ketogenic diet slows down mitochondrial myopathy progression in mice. Hum Mol Genet 19: 1974–1984

Anderson NS, Haynes CM (2020) Folding the Mitochondrial UPR into the Integrated Stress Response. Trends Cell Biol 30: 428–439

Becker C, Kukat A, Szczepanowska K, Hermans S, Senft K, Brandscheid CP, Maiti P, Trifunovic A (2018) CLPP deficiency protects against metabolic syndrome but hinders adaptive thermogenesis. EMBO Rep 19

Bhaskaran S, Pharaoh G, Ranjit R, Murphy A, Matsuzaki S, Nair BC, Forbes B, Gispert S, Auburger G, Humphries KM et al (2018) Loss of mitochondrial protease ClpP protects mice from diet-induced obesity and insulin resistance. EMBO Rep 19

Canto C, Garcia-Roves PM (2015) High-Resolution Respirometry for Mitochondrial Characterization of Ex Vivo Mouse Tissues. Curr Protoc Mouse Biol 5: 135–153

Chacinska A, Koehler CM, Milenkovic D, Lithgow T, Pfanner N (2009) Importing mitochondrial proteins: machineries and mechanisms. Cell 138: 628–644

Chacinska A, Lind M, Frazier AE, Dudek J, Meisinger C, Geissler A, Sickmann A, Meyer HE, Truscott KN, Guiard B et al (2005) Mitochondrial presequence translocase: switching between TOM tethering and motor recruitment involves Tim21 and Tim17. Cell 120: 817–829

Choi MS, Kim YJ, Kwon EY, Ryoo JY, Kim SR, Jung UJ (2015) High-fat diet decreases energy expenditure and expression of genes controlling lipid metabolism, mitochondrial function and skeletal system development in the adipose tissue, along with increased expression of extracellular matrix remodelling- and inflammation-related genes. Br J Nutr 113: 867–877

Coll AP, Chen M, Taskar P, Rimmington D, Patel S, Tadross JA (2020) GDF15 mediates the effects of metformin on body weight and energy balance. Nature 578: 444–448

Dobin A, Davis CA, Schlesinger F, Drenkow J, Zaleski C, Jha S, Batut P, Chaisson M, Gingeras TR (2013) STAR: ultrafast universal RNA-seq aligner. Bioinformatics 29: 15–21

Durieux J, Wolff S, Dillin A (2011) The cell-non-autonomous nature of electron transport chain-mediated longevity. Cell 144: 79–91

Fabregat A, Sidiropoulos K, Garapati P, Gillespie M, Hausmann K, Haw R, Jassal B, Jupe S, Korninger F, McKay S et al (2016) The Reactome pathway Knowledgebase. Nucleic Acids Research 44: D481–D487

Forsstrom S, Jackson CB, Carroll CJ, Kuronen M, Pirinen E, Pradhan S, Marmyleva A, Auranen M, Kleine IM, Khan NA et al (2019) Fibroblast Growth Factor 21 Drives Dynamics of Local and Systemic Stress Responses in Mitochondrial Myopathy with mtDNA Deletions. Cell Metab 30: 1040–1054 e1047

Friedman JR, Nunnari J (2014) Mitochondrial form and function. Nature 505: 335–343

Heinonen S, Buzkova J, Muniandy M, Kaksonen R, Ollikainen M, Ismail K, Hakkarainen A, Lundbom J, Lundbom N, Vuolteenaho K et al (2015) Impaired Mitochondrial Biogenesis in Adipose Tissue in Acquired Obesity. Diabetes 64: 3135–3145

Heinonen S, Muniandy M, Buzkova J, Mardinoglu A, Rodriguez A, Fruhbeck G, Hakkarainen A, Lundbom J, Lundbom N, Kaprio J et al (2017) Mitochondria-related transcriptional signature is downregulated in adipocytes in obesity: a study of young healthy MZ twins. Diabetologia 60: 169–181

Horst M, Oppliger W, Rospert S, Schonfeld HJ, Schatz G, Azem A (1997) Sequential action of two hsp70 complexes during protein import into mitochondria. EMBO J 16: 1842–1849

Jiang S, Yuan T, Rosenberger FA, Mourier A, Dragano NRV, Kremer LS, Rubalcava-Gracia D, Hansen FM, Borg M, Mennuni M et al (2024) Inhibition of mammalian mtDNA transcription acts paradoxically to reverse diet-induced hepatosteatosis and obesity. Nat Metab 6: 1024–1035

Kampinga HH, Craig EA (2010) The HSP70 chaperone machinery: J proteins as drivers of functional specificity. Nat Rev Mol Cell Biol 11: 579–592

Kang PJ, Ostermann J, Shilling J, Neupert W, Craig EA, Pfanner N (1990) Requirement for hsp70 in the mitochondrial matrix for translocation and folding of precursor proteins. Nature 348: 137–143

Khan NA, Nikkanen J, Yatsuga S, Jackson C, Wang L, Pradhan S, Kivela R, Pessia A, Velagapudi V, Suomalainen A (2017) mTORC1 Regulates Mitochondrial Integrated Stress Response and Mitochondrial Myopathy Progression. Cell Metab 26: 419–428 e415

Konovalova S, Liu X, Manjunath P, Baral S, Neupane N, Hilander T, Yang Y, Balboa D, Terzioglu M, Euro L et al (2018) Redox regulation of GRPEL2 nucleotide exchange factor for mitochondrial HSP70 chaperone. Redox Biol 19: 37–45

Krayl M, Lim JH, Martin F, Guiard B, Voos W (2007) A cooperative action of the ATP-dependent import motor complex and the inner membrane potential drives mitochondrial preprotein import. Mol Cell Biol 27: 411–425

Lallemand Y, Luria V, Haffner-Krausz R, Lonai P (1998) Maternally expressed PGK-Cre transgene as a tool for early and uniform activation of the Cre site-specific recombinase. Transgenic Res 7: 105–112

Liao Y, Smyth GK, Shi W (2014) featureCounts: an efficient general purpose program for assigning sequence reads to genomic features. Bioinformatics 30: 923–930

Lill R, Freibert SA (2020) Mechanisms of Mitochondrial Iron-Sulfur Protein Biogenesis. Annu Rev Biochem 89: 471–499

Lin YF, Schulz AM, Pellegrino MW, Lu Y, Shaham S, Haynes CM (2016) Maintenance and propagation of a deleterious mitochondrial genome by the mitochondrial unfolded protein response. Nature 533: 416–419

Liu Q, D’Silva P, Walter W, Marszalek J, Craig EA (2003) Regulated cycling of mitochondrial Hsp70 at the protein import channel. Science 300: 139–141

Love MI, Huber W, Anders S (2014) Moderated estimation of fold change and dispersion for RNA-seq data with DESeq2. Genome Biology 15: 550–550

Manjunath P, Stojkovic G, Euro L, Konovalova S, Wanrooij S, Koski K, Tyynismaa H (2024) Preferential binding of ADP-bound mitochondrial HSP70 to the nucleotide exchange factor GRPEL1 over GRPEL2. Protein Sci 33: e5190

Milenkovic D, Ramming T, Muller JM, Wenz LS, Gebert N, Schulze-Specking A, Stojanovski D, Rospert S, Chacinska A (2009) Identification of the signal directing Tim9 and Tim10 into the intermembrane space of mitochondria. Mol Biol Cell 20: 2530–2539

Model K, Meisinger C, Prinz T, Wiedemann N, Truscott KN, Pfanner N, Ryan MT (2001) Multistep assembly of the protein import channel of the mitochondrial outer membrane. Nat Struct Biol 8: 361–370

Morizono MA, McGuire KL, Birouty NI, Herzik MA, Jr. (2024) Structural insights into GrpEL1-mediated nucleotide and substrate release of human mitochondrial Hsp70. Nat Commun 15: 10815

Morizono MA, Safar TV, Herzik MA (2026) Tuning the Hsp70 chaperone cycle: emerging roles of GrpE-like nucleotide exchange factors in proteostasis and organelle function. Protein Cell 17: 176–189

Muller TD, Klingenspor M, Tschop MH (2021) Revisiting energy expenditure: how to correct mouse metabolic rate for body mass. Nat Metab 3: 1134–1136

Naylor DJ, Stines AP, Hoogenraad NJ, Hoj PB (1998) Evidence for the existence of distinct mammalian cytosolic, microsomal, and two mitochondrial GrpE-like proteins, the Co-chaperones of specific Hsp70 members. J Biol Chem 273: 21169–21177

Neupane N, Rajendran J, Kvist J, Harjuhaahto S, Hu B, Kinnunen V, Yang Y, Nieminen AI, Tyynismaa H (2022) Inter-organellar and systemic responses to impaired mitochondrial matrix protein import in skeletal muscle. Commun Biol 5: 1060

Neupert W, Herrmann JM (2007) Translocation of proteins into mitochondria. Annu Rev Biochem 76: 723–749

Ott M, Amunts A, Brown A (2016) Organization and Regulation of Mitochondrial Protein Synthesis. Annu Rev Biochem 85: 77–101

Pagliarini DJ, Calvo SE, Chang B, Sheth SA, Vafai SB, Ong SE, Walford GA, Sugiana C, Boneh A, Chen WK et al (2008) A mitochondrial protein compendium elucidates complex I disease biology. Cell 134: 112–123

Quiros PM, Prado MA, Zamboni N, D’Amico D, Williams RW, Finley D, Gygi SP, Auwerx J (2017) Multi-omics analysis identifies ATF4 as a key regulator of the mitochondrial stress response in mammals. J Cell Biol 216: 2027–2045

Rajendran J, Purhonen J, Tegelberg S, Smolander OP, Morgelin M, Rozman J, Gailus-Durner V, Fuchs H, Hrabe de Angelis M, Auvinen P et al (2019) Alternative oxidase-mediated respiration prevents lethal mitochondrial cardiomyopathy. EMBO Mol Med 11

Saitoh T, Igura M, Obita T, Ose T, Kojima R, Maenaka K, Endo T, Kohda D (2007) Tom20 recognizes mitochondrial presequences through dynamic equilibrium among multiple bound states. EMBO J 26: 4777–4787

Slutsky-Leiderman O, Marom M, Iosefson O, Levy R, Maoz S, Azem A (2007) The interplay between components of the mitochondrial protein translocation motor studied using purified components. J Biol Chem 282: 33935–33942

Srivastava S, Savanur MA, Sinha D, Birje A, R V, Saha PP, D’Silva P (2017) Regulation of mitochondrial protein import by the nucleotide exchange factors GrpEL1 and GrpEL2 in human cells. J Biol Chem 292: 18075–18090

Suomalainen A, Battersby BJ (2018) Mitochondrial diseases: the contribution of organelle stress responses to pathology. Nat Rev Mol Cell Biol 19: 77–92

Suomalainen A, Nunnari J (2024) Mitochondria at the crossroads of health and disease. Cell 187: 2601–2627

Team RC, 2019. R: A Language and Environment for Statistical Computing. Vienna, Austria.

Templeman NM, Flibotte S, Chik JHL, Sinha S, Lim GE, Foster LJ, Nislow C, Johnson JD (2017) Reduced Circulating Insulin Enhances Insulin Sensitivity in Old Mice and Extends Lifespan. Cell Rep 20: 451–463

Tschop MH, Speakman JR, Arch JR, Auwerx J, Bruning JC, Chan L, Eckel RH, Farese RV, Jr., Galgani JE, Hambly C et al (2011) A guide to analysis of mouse energy metabolism. Nat Methods 9: 57–63

Tyynismaa H, Mjosund KP, Wanrooij S, Lappalainen I, Ylikallio E, Jalanko A, Spelbrink JN, Paetau A, Suomalainen A (2005) Mutant mitochondrial helicase Twinkle causes multiple mtDNA deletions and a late-onset mitochondrial disease in mice. Proc Natl Acad Sci U S A 102: 17687–17692

van der Laan M, Meinecke M, Dudek J, Hutu DP, Lind M, Perschil I, Guiard B, Wagner R, Pfanner N, Rehling P (2007) Motor-free mitochondrial presequence translocase drives membrane integration of preproteins. Nat Cell Biol 9: 1152–1159

van der Leij FR, Bloks VW, Grefhorst A, Hoekstra J, Gerding A, Kooi K, Gerbens F, te Meerman G, Kuipers F (2007) Gene expression profiling in livers of mice after acute inhibition of beta-oxidation. Genomics 90: 680–689

Vernochet C, Mourier A, Bezy O, Macotela Y, Boucher J, Rardin MJ, An D, Lee KY, Ilkayeva OR, Zingaretti CM et al (2012) Adipose-specific deletion of TFAM increases mitochondrial oxidation and protects mice against obesity and insulin resistance. Cell Metab 16: 765–776

Wiedemann N, Pfanner N (2017) Mitochondrial Machineries for Protein Import and Assembly. Annu Rev Biochem 86: 685–714

Wrobel L, Topf U, Bragoszewski P, Wiese S, Sztolsztener ME, Oeljeklaus S, Varabyova A, Lirski M, Chroscicki P, Mroczek S et al (2015) Mistargeted mitochondrial proteins activate a proteostatic response in the cytosol. Nature 524: 485–488

Yu G, He Q-Y (2016) ReactomePA: an R/Bioconductor package for reactome pathway analysis and visualization. Mol BioSyst 12: 477–479

Yu G, Wang L-G, Han Y, He Q-Y (2012) clusterProfiler: an R Package for Comparing Biological Themes Among Gene Clusters. OMICS : a Journal of Integrative Biology 16: 284–287

