## Supplementary Figures for "Loss of mitochondrial co-chaperone GRPEL2 protects mice from age- and diet-induced obesity"

**Supplementary Figure 1**

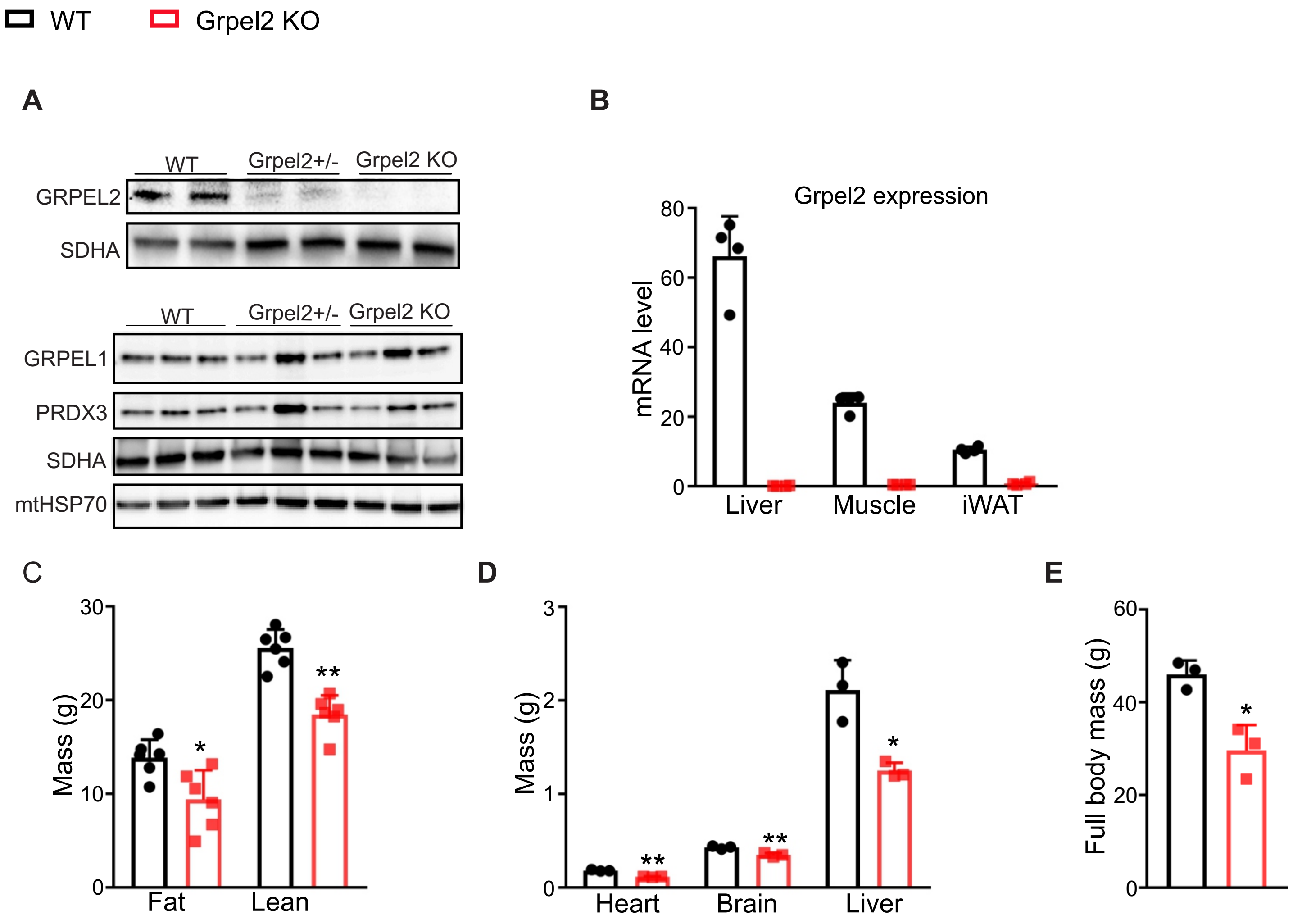

### Supplementary Figure 2

WT    Grpel2 KO

**1y old**

**A**

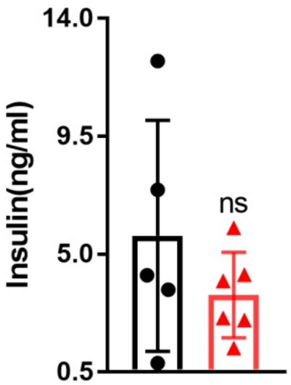


**High fat diet (HFD)**

**B**

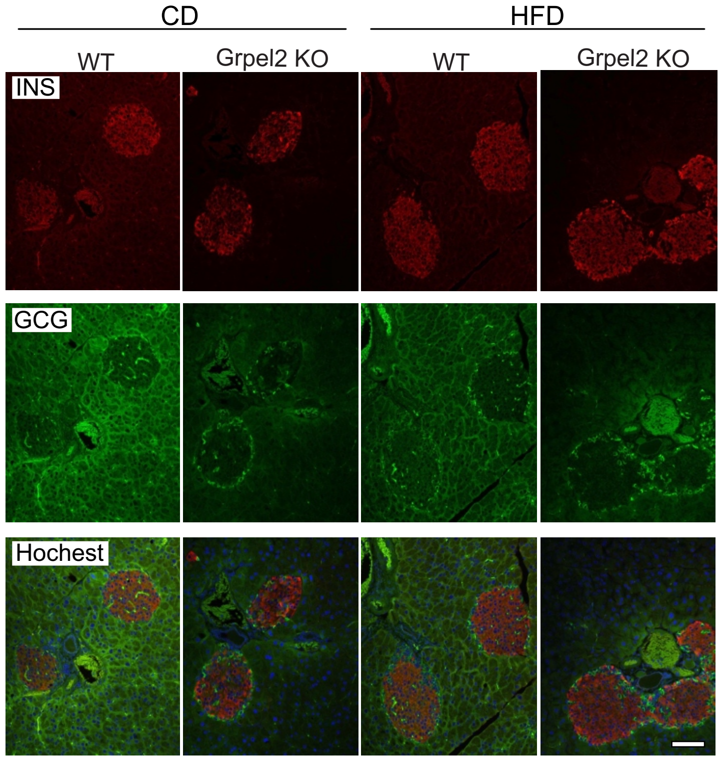


**C**

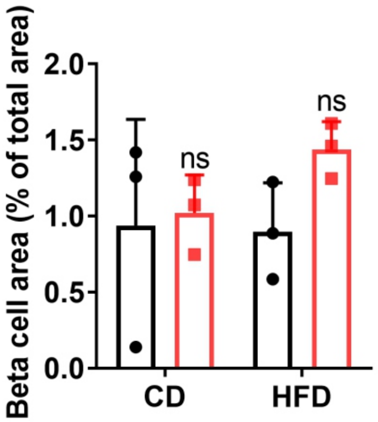

**Supplementary Figure 3**

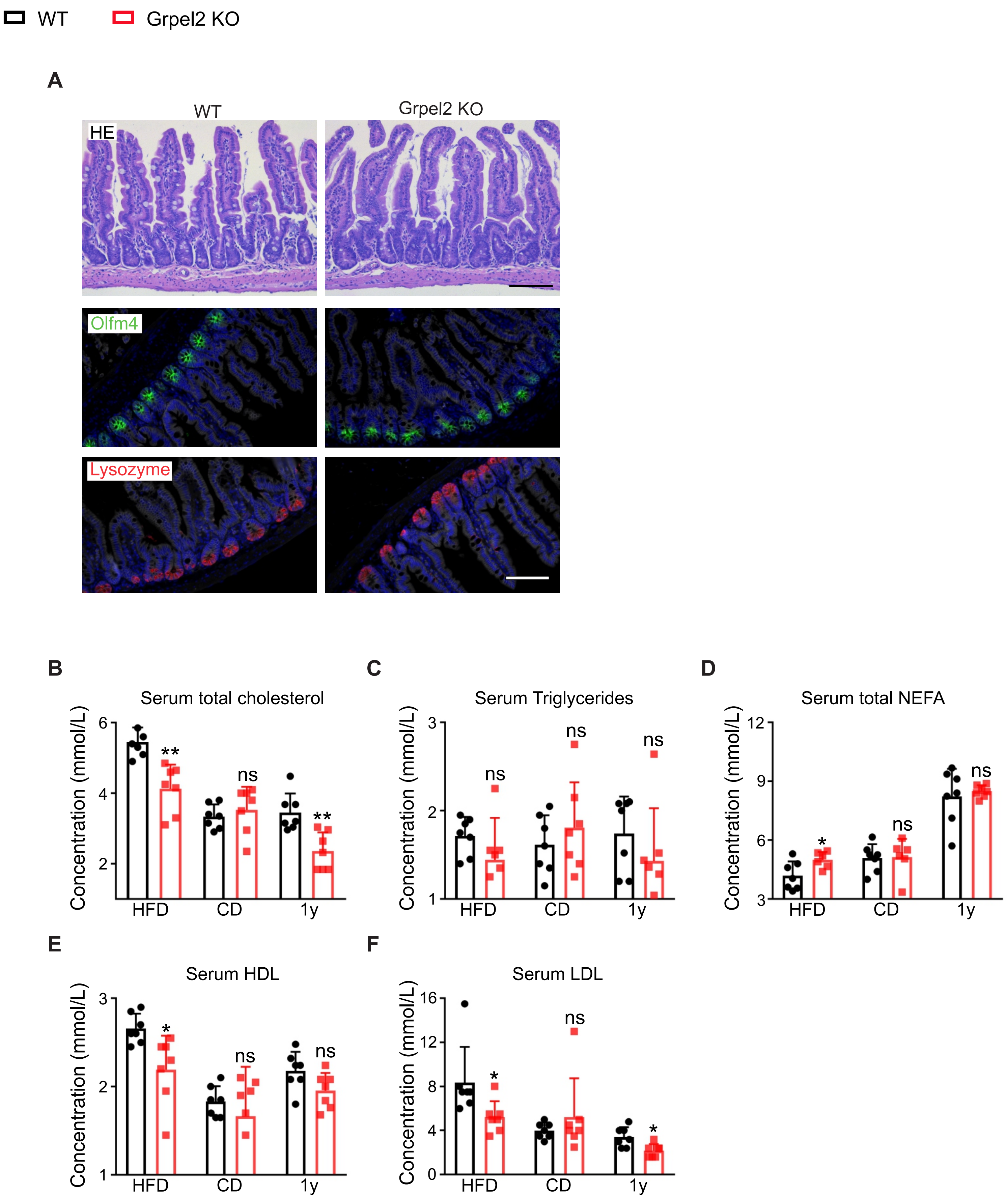

### Supplementary Figure 4

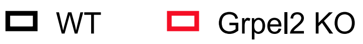


**A**

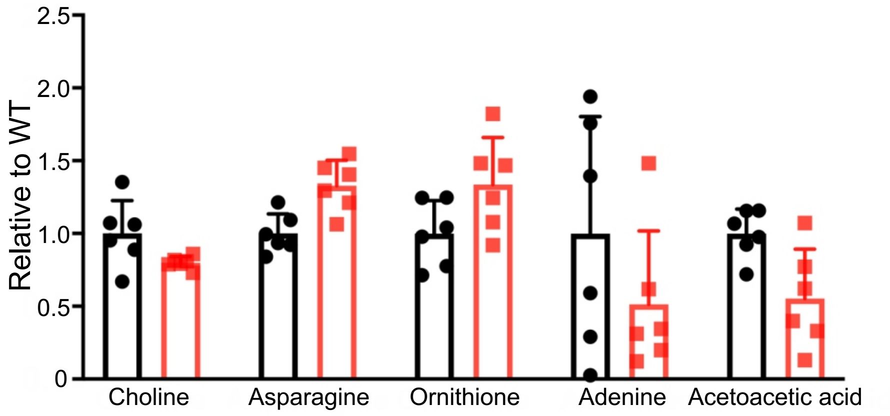


**B**

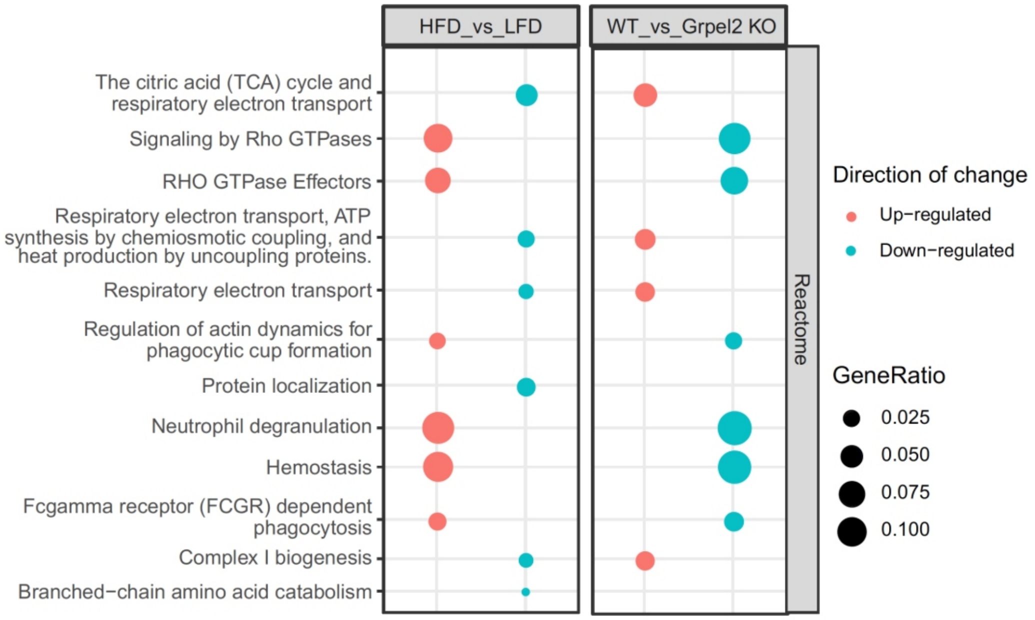
